# Identification of bipotent progenitors that give rise to myogenic and connective tissues in mouse

**DOI:** 10.1101/2021.05.26.445757

**Authors:** Alexandre Grimaldi, Glenda Comai, Sébastien Mella, Shahragim Tajbakhsh

**Affiliations:** Stem Cells & Development Unit, 25 rue du Dr. Roux, Institut Pasteur, 75015 Paris, France; UMR CNRS 3738, Institut Pasteur, Paris, France

## Abstract

How distinct cell fates are manifested by direct lineage ancestry from bipotent progenitors, or by specification of individual cell types within a field of cells is a key question for understanding the emergence of tissues. The interplay between skeletal muscle progenitors and associated connective tissues cells provides a model for examining how muscle functional units are established. Most craniofacial structures originate from the vertebrate-specific neural crest cells except in the dorsal portion of the head, where they arise from cranial mesoderm. Here, using multiple lineage-traced single cell RNAseq, advanced computational methods and in situ analyses, we identify Myf5^+^ bipotent progenitors that give rise to both muscle and juxtaposed connective tissue. Following this bifurcation, muscle and connective tissue cells retain complementary signalling features and maintain spatial proximity. Interruption of upstream myogenic identity shifts muscle progenitors to a connective tissue fate. Interestingly, Myf5-derived connective tissue cells, which adopt a novel regulatory signature, were not observed in ventral craniofacial structures that are colonised by neural crest cells. Therefore, we propose that an ancestral program gives rise to bifated muscle and connective tissue cells in skeletal muscles that are deprived of neural crest.

## INTRODUCTION

Stromal cells that are associated with skeletal muscles play critical roles in providing structural support and molecular cues (Biferali et al., 2019; Kardon et al., 2003; Sefton and Kardon, 2019). The majority of muscle-associated connective tissues in the head is derived from cranial neural crest cells (NCCs), an embryonic cell population that contributes to most of the structural components of the “new head”, a vertebrate innovation (Douarin and Kalcheim, 1999; Gans and Northcutt, 1983; Grenier et al., 2009; Heude et al., 2018; Noden and Trainor, 2005). Recently, the extent of this contribution was redefined in muscles derived from cranial mesoderm, including extraocular (EOM), laryngeal and pharyngeal muscles (Comai et al., 2020; Grimaldi et al., 2015; Heude et al., 2018; Noden and Epstein, 2010). Interestingly, these muscles contain mesenchyme that is mesoderm-derived in their dorso-medial component, whereas the remaining muscle mass is embedded in mesenchyme that is neural crest-derived. It is unclear how the coordinated emergence of myogenic and connective tissues takes place during development, and how they establish long lasting paracrine communication.

Along the trunk axis, paraxial somitic mesoderm gives rise to skeletal muscles and associated connective tissues (Burke and Nowicki, 2003). Upon signals emanating from the neural tube, notochord, ectoderm and lateral plate mesoderm, the dermomyotome (dorsal portion of the somite) undergoes an epithelial-to-mesenchymal transition and gives rise to several cell types including all skeletal muscles of the body, vasculature, tendons and bones (Ben-Yair and Kalcheim, 2008; Christ et al., 2007). Similarly, cranial mesodermal progenitors give rise to these diverse cell types yet, the unsegmented nature of head mesoderm raises the question of how spatiotemporal control of these cellular identities is established. Moreover, cardiopharyngeal mesoderm, which constitutes the major portion of cranial mesoderm, has cardiovascular potential, which manifests in the embryo as regions of clonally related cardiac and craniofacial skeletal muscles (Diogo et al., 2015; Swedlund and Lescroart, 2019). This skeletal muscle/cardiac branchpoint has been the subject of intense investigation in several model organisms including ascidians, avians, and mouse (Wang et al., 2019). However, the issue of connective tissue divergence from this or another lineage has not been addressed.

Recently, advanced pipelines integrating scRNAseq data and modern algorithms have been instrumental for identifying new lineage relationships during development (Cao et al., 2019; He et al., 2020). Here, we employed unbiased and lineage-restricted single-cell transcriptomics using multiple transgenic mouse lines combined with new computational methods, in situ labelling and loss of function experiments, and show that bipotent progenitors expressing the muscle determination gene *Myf5* give rise to skeletal muscle and anatomically associated connective tissues. Notably, this property was restricted to muscles with partial contribution from NCCs, suggesting that in the absence of NCCs, cranial mesoderm acts as a source of connective tissue.

## RESULTS

### Myogenic and non-myogenic mesodermal populations coexist within distinct head lineages

Somitic (*Pax3*-dependent) and cranial (*Tbx1/Pitx2-dependent*) mesoderm give rise to diverse cell types including those of the musculoskeletal system (Figure 1A). We first set out to explore the emergence of skeletal muscles and their associated mesodermal tissue within these programs. To that end, we employed a broad anterior mesoderm lineage-tracing strategy using *Mesp1^Cre/+^;R26^mTmG/+^* mouse embryos at E10.5 when craniofacial skeletal muscles are being established (Heude et al., 2018). The upper third (anterior to forelimb) of the embryos was dissected, the live GFP+ cells were isolated by FACS and processed for scRNAseq analysis (Figure S1A, C-D). After removal of doublets and lower quality cells (see Methods), a large portion of the cells obtained corresponded to adipogenic, chondrogenic, sclerotomal, endothelial, and cardiovascular cells (Figure 1B, Figure S2A-B). *Pax3*, *Pitx2*, *Tbx1*, *Myf5* and *Myod* expression distinguished clusters containing the cranial myogenic progenitors (Figure1C, Figure S2A).

**Figure 1.**
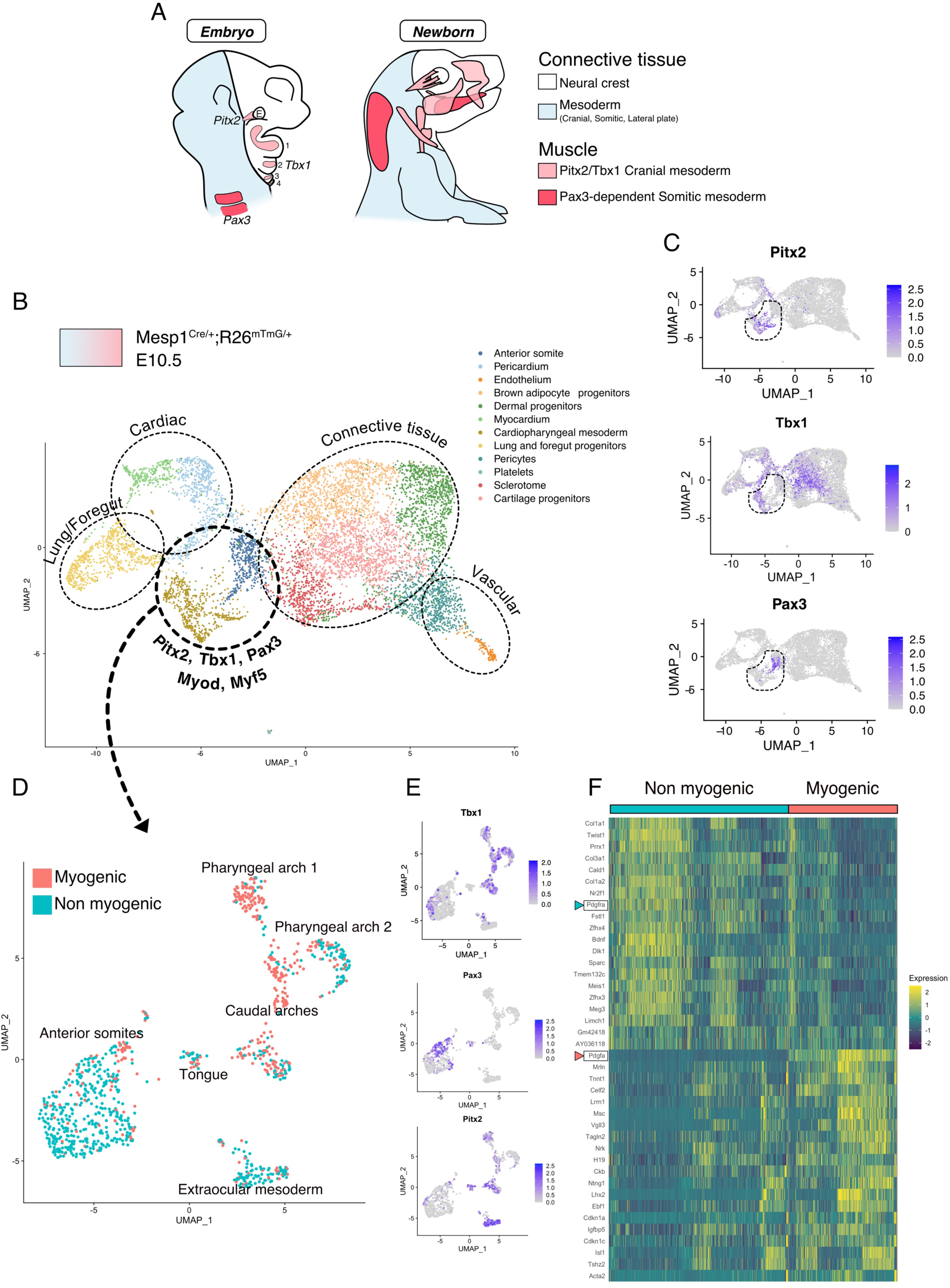
scRNAseq reveals non-myogenic populations of cranial mesoderm lineages. A) Scheme of connective tissue origin in the head and known mesodermal upstream regulators. E: Eye, 1-4: Pharyngeal arches 1-4. B) UMAP of *Mesp1^Cre/+^; R26^mTmG/+^* E10.5 scRNAseq with main cell types highlighted. C) UMAP expression plots of *Pitx2* (EOM), *Tbx1* (cranial mesoderm except EOM) and *Pax3* (somitic mesoderm), indicating the clusters of progenitors. D) UMAP of progenitor subset annotated as myogenic and non-myogenic based on expression patterns found in E and F. E) UMAP expression plots *Pitx2*, *Tbx1* and *Pax3*. F) Heatmap of top 20 markers of myogenic versus non-myogenic clusters. *Pdgfra/Pdgfa* genes are highlighted.

After subsetting these clusters, a few subclusters clearly separated based on their origin and anatomical location (Figure1D-E, Figure S2C). Surprisingly, about half of the cells exhibited a connective tissue signature, including a strong bias towards *Prrx1*, a marker for lateral plate mesoderm (Durland et al., 2008), *Col1a1*, a major extracellular matrix component of connective cells (Micheli et al., 2020), and *Twist1*, previously reported to confer mesenchymal properties to cranial mesoderm (Bildsoe et al., 2016) (Figure 1F). Furthermore, the expression of *Pdgfra*, a well-defined marker of stromal cells (Farahani and Xaymardan, 2015), was robustly anticorrelated with the expression of its ligand *Pdgfa*, and associated with non-myogenic genes. Conversely, *Pdgfa* expression correlated with a myogenic cell state (Figure1F, Figure S2D). Myogenic *Pdgfa* expression was shown to promote adjacent sclerotomal cells to adopt a rib cartilage fate (Tallquist et al., 2000). Therefore, this analysis identified anatomically distinct muscle and closely-associated connective tissue progenitors and highlights a potential PDGFR-mediated crosstalk between these 2 cells types.

### Transcriptional trajectories reveal a myogenic to non-myogenic cell state transition

To understand the lineage relationship between myogenic and non-myogenic cells, we exploited the unspliced and spliced variants of our scRNAseq data, and computed the RNA velocity in each cell, using a recently described tool (Bergen et al., 2020) (Figure 2, Figure S3). RNA velocity interrogates the relative abundance of unspliced and spliced gene variants, which depends on the rates of transcription, degradation, and splicing to infer directional trajectories (Bergen et al., 2020; Manno et al., 2018). The cell cycle status constitutes a potential bias in scRNAseq data, especially when heterogeneous populations undergo cellular expansion, commitment and differentiation (McDavid et al., 2016). To eliminate this potential bias, cell cycle genes were consistently regressed out during preprocessing and directional trajectories were overlaid with cell cycle phase visualization for comparisons (Figure S3A, Methods). Notably, RNA velocity-inferred trajectories indicated that Myf5+ cells from the myogenic compartment contributed to non-myogenic cells (Figure 2A). These calculations were based on gene- and cluster-specific dynamics, which yield higher accuracy than the initially described RNA velocity method, while providing quantitative metrics for quality control (Figure S3B and Methods).

**Figure 2.**
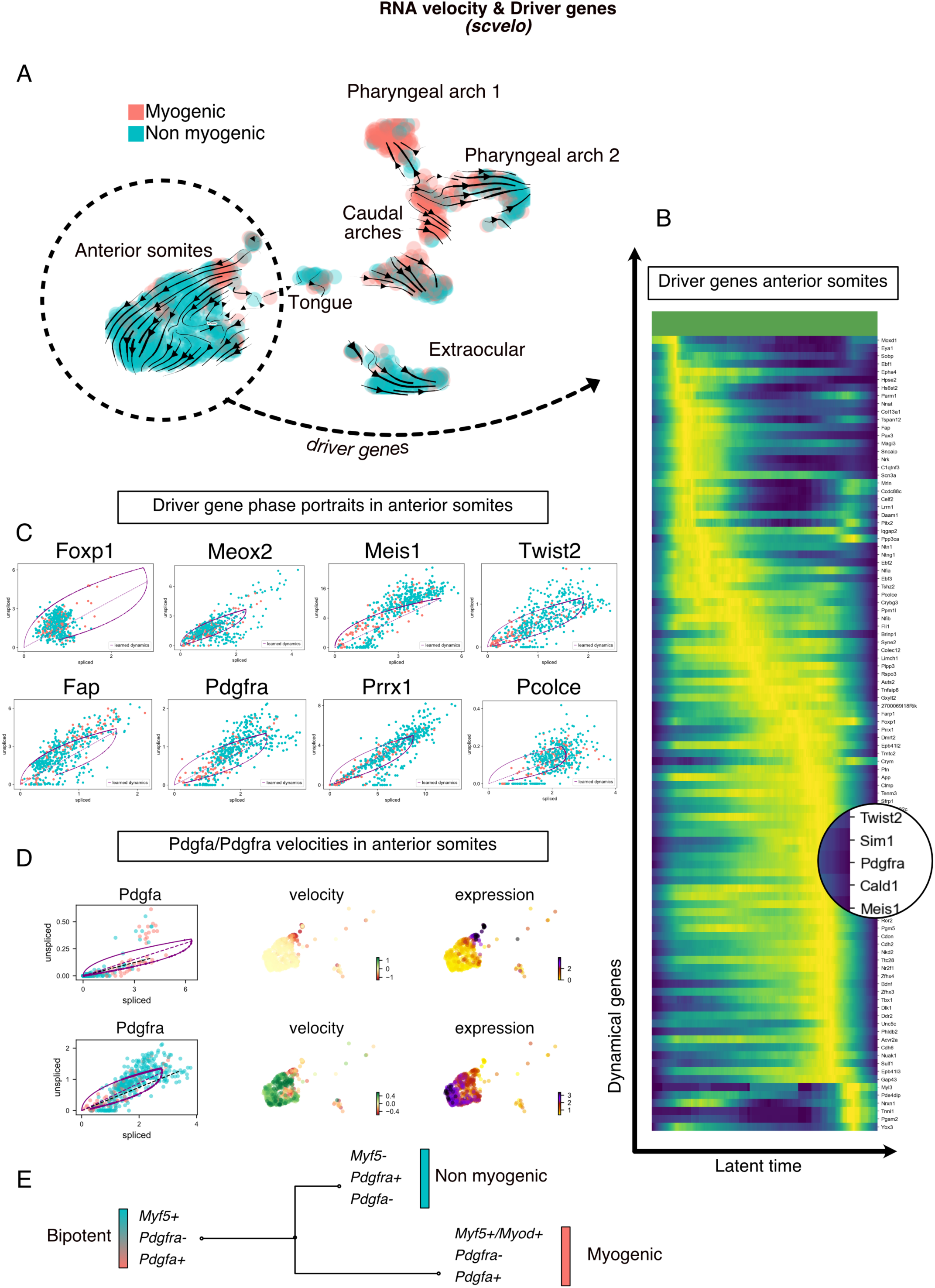
Transcriptomic dynamics reveal a myogenic to non-myogenic transition in anterior somite progenitors. A) Velocity UMAP plots displaying myogenic and non-myogenic clusters. Arrows represent the lineage progression based on RNA velocity (relative abundance of unspliced and spliced transcripts). B) Heatmap of driver genes accounting for anterior somite velocity, highlighting *Pdgfra*. C) Phase portraits of few selected driver genes in the anterior somites: *Foxp1, Meox2, Meis1, Twist2, Fap, Pdgfra, Prrx1* and *Pcolce*. Y-axis represents the amount of unspliced transcript per cell; X-axis represents the number of spliced transcripts per cell. A high fraction of unspliced variants indicates an active transcription of the locus, while the inverse indicates inactive/repressed transcription. Dynamics of transcription were inferred at a gene- and cluster-specific level (see Methods). D) Phase portraits, velocity and expression plots of *Pdgfa* and *Pdgfra* showing splicing dynamics of these 2 genes. E) Working model of myogenic and non-myogenic fate decision from a common bipotent progenitor in anterior somites.

Another powerful feature of this method is the ability to infer “driver genes” that are responsible for most of the calculated RNA velocity, hence actively transcribed, or repressed (Bergen et al., 2020). Therefore, these genes can identify transitory states underlying cell fate decisions. We used this approach to uncover the driver genes that were responsible for the velocity found in anterior somites, as these cells displayed the most consistent directionality, and appeared to be independent of cell cycle (Figure 2B, Figure S3A-B, Table1). Top transcribed driver genes included *Foxp1* (Shao and Wei, 2018)*, Meox2* (Noizet et al., 2016)*, Meis1* (López-Delgado et al., 2020)*, Twist2* (Franco et al., 2009) *, Fap* (Puré and Blomberg, 2018)*, Pdgfra* (Tallquist et al., 2000)*, Prrx1* (Leavitt et al., 2020) and *Pcolce* (Bildsoe et al., 2016), that are associated with fibrosis and connective tissue development (Figure 2C). Interestingly, we noted that *Pdgfra* appeared as a driver gene and was activated along this inferred trajectory, whereas *Pdgfa* expression decreased rapidly (Figure 2D). Taken together, RNA velocity analysis for anterior somite mesodermal progenitors showed that Myf5+/Pdgfa+ cells shifted towards a non-myogenic fate, by downregulating these 2 markers and activating *Pdgfra* expression (Figure 2E).

**Table 1:**
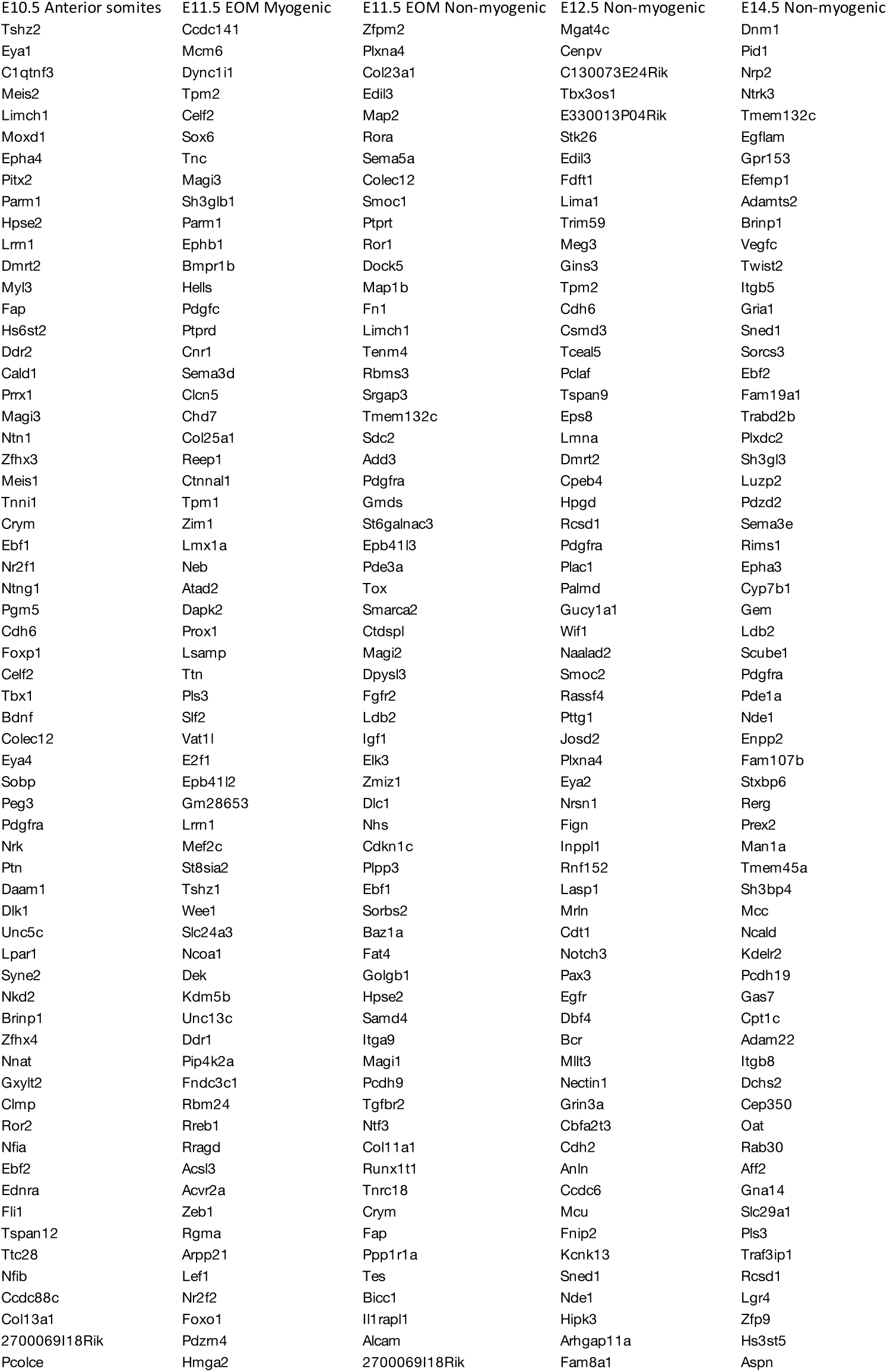
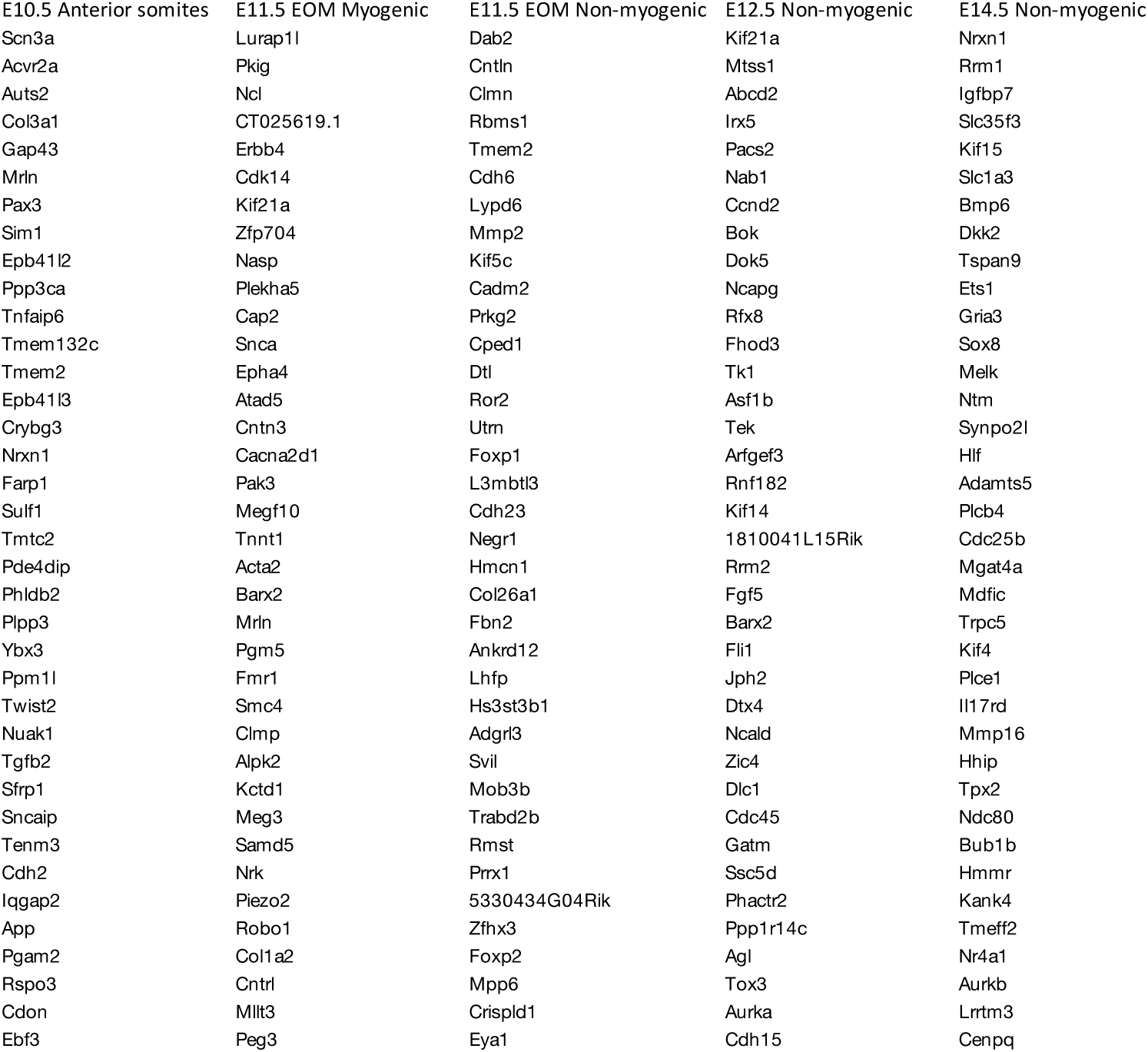
Driver genes underlying cell fate decision in each dataset.

### *Myf5*-derived lineage contributes to connective tissue cells in the absence of neural crest

Given that the number of cells examined in the EOM and pharyngeal arch mesodermal clusters from the E10.5 dataset was lower than for anterior somites, we decided to validate the relevance of Myf5-derived non-myogenic cells in these cranial regions directly in vivo. We thus examined the EOM, larynx and upper back muscles in the early fetus at E14.5 using a *Myf5*-lineage reporter mouse (*Myf5^Cre/+^; R26^TdTomato/+^*) combined with a contemporary reporter for Pdgfra+ non-myogenic cells (*Pdgfra^H2BGFP/+^*) (Figure 3). Notably, we observed tdTom/H2BGFP double-positive cells in regions of EOM, laryngeal and upper back muscles that were partially or fully deprived of neural crest (Adachi et al., 2020; Comai et al., 2020; Heude et al., 2018) (Figure 3A-C’). Conversely, no double-positive cells were detected in muscles that are fully embedded in neural crest derived connective tissues such as mandibular and tongue muscles (Heude et al., 2018) (Figure 3D-E’).

**Figure 3.**
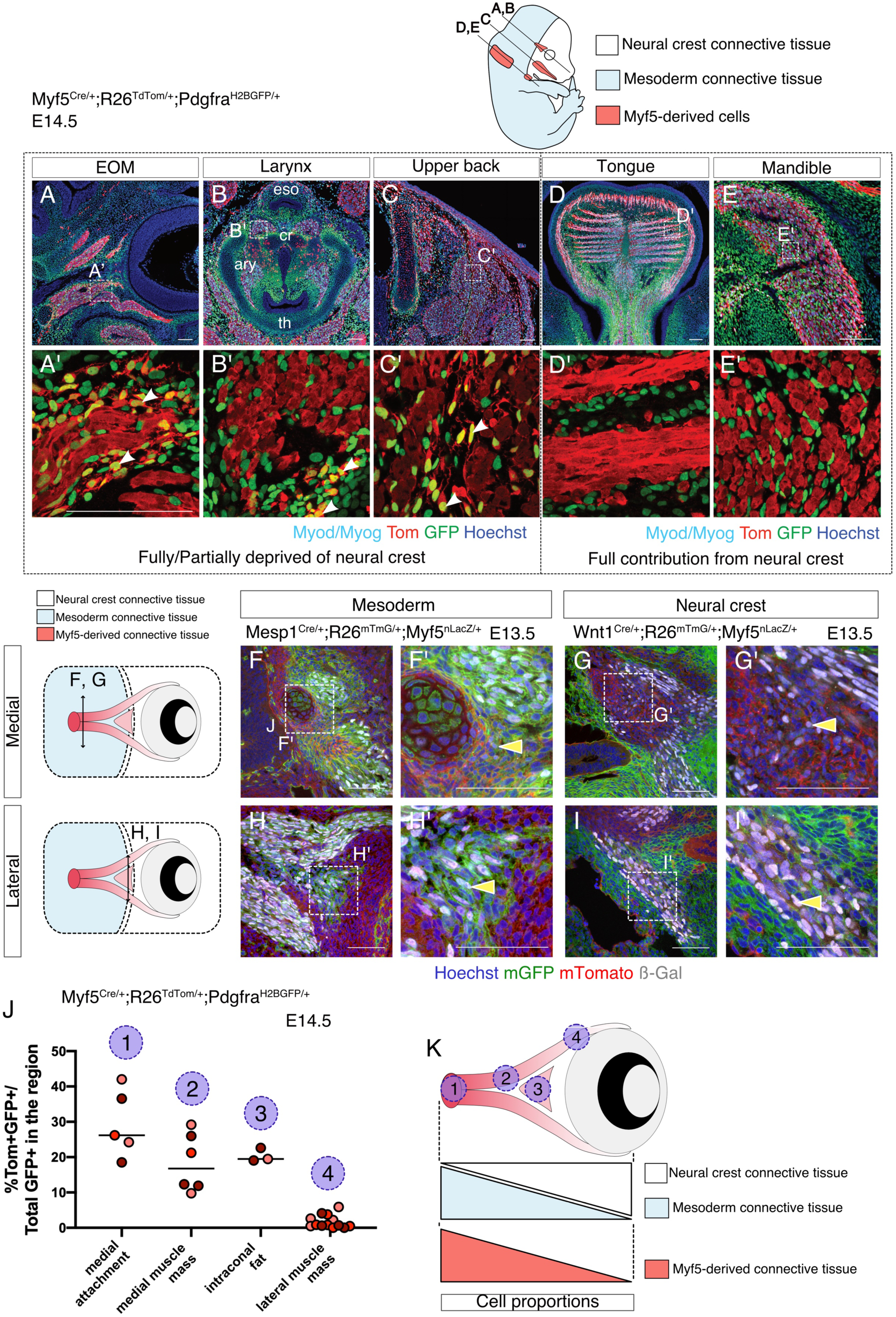
Myf5-derived mesodermal connective tissue partially compensates for the lack of neural crest. A-E’) Transverse sections of an E14.5 *Myf5^Cre/+^; R26^TdTomato/+^; Pdgfra^H2BGFP/+^* embryo immunostained for Myod/Myog. White arrowheads indicated cells double-positive GFP/Tomato and negative for Myod/Myog (n=3 embryos). F-I’) Transverse cryosections of the EOM at E13.5 of *Wnt1^Cre/+^; R26^mTmG/+^; Myf5^nlacZ/+^* (G,I) and *Mesp1^Cre/+^; R26^mTmG/+^; Myf5^nlacZ/+^* (F,H) immunostained for β-gal, at the level of the medial attachment (F,G) and lateral muscle masses (H,I). Yellow arrowhead indicates *Myf5*-expressing cells in the context of mesodermal and neural crest lineages. (n=2 embryos with 5 tissue sections analyzed per embryo) J) Quantifications of the proportion of double positive cells in E14.5 *Myf5^Cre/+^; R26^TdTomato/+^; Pdgfra^H2BGFP/+^* embryo in various regions throughout the EOM (n=3 embryos, with 5 tissue sections analyzed per embryo). K) Scheme highlighting the quantified regions in (J), and summarising the contribution of each population to connective tissue.

Mesenchymal tissue that is associated with the EOM arises from mesoderm in its most dorso-medial portion and from neural crest in its ventro-lateral portion (Comai et al., 2020). This dual origin makes it a prime candidate to explore the relative contribution of *Myf5*-derived cells to the associated connective tissues within a single functional unit. Using *Wnt1^Cre/+^; R26^mTmG/+^; Myf5^nlacZ/+^* (NCC tracing) and *Mesp1^Cre/+^;R26^mTmG/+^;Myf5^nlacZ/+^* (mesoderm tracing) at E13.5, we found that *Myf5*-expressing cells (assessed by β-gal expression) were exclusively in *Mesp1*-derived domains and absent from the *Wnt1* lineage (Figure 3F-I’). By E14.5, we observed a medio-lateral gradient of *Myf5*-lineage contribution to EOM associated connective tissues, and this was anticorrelated with the local contribution of neural crest cells to connective tissues (Figure 3J-K).

In agreement with our scRNAseq velocity analysis, these observations suggest that the mesodermal *Myf5*-lineage compensates for the lack of muscle-associated connective tissue in domains that are deprived of neural crest.

### Myf5-derived cells can maintain a molecular crosstalk following bifurcation in cell fate

To investigate in more detail potential paracrine cell-cell communication between myogenic and non-myogenic cells, we examined their signalling complementarity together with their anatomical proximity. We first generated an E11.5 *Myf5*-lineage traced (*Myf5^Cre/+^;R26^mTmG/+^*) sc-RNAseq dataset (Figure S1A, E-F, Figure S3A, C) and focused on the EOM, which was clearly distinguished as an independent cluster based on the co-expression of several markers including *Pitx2* and *Alx4* (Bothe and Dietrich, 2006) (Figure 4A). Here, RNA velocity revealed strong myogenic/non-myogenic bi-directional cell-fate transitions (Figure 4B). In agreement with the E10.5 mesodermal sc-RNAseq dataset, EOM progenitors presented a strong dichotomy in *Pdgfa* and *Pdfgra* expression between myogenic and non-myogenic cells, respectively. To confirm if the potential crosstalk between these clusters is maintained at later stages of EOM development, we interrogated the anatomical proximity of these cells by performing single molecule fluorescent in situ hybridization (RNAscope) for *Pdgfa* and *Pdgfra* on E14.5 lineage-traced *Myf5^Cre/+^;R26^mTmG/+^* fetuses (Figure 4C-D’). In accordance with the scRNAseq analysis, we observed a complementary pattern of expression of *Pdgfa* (membrane GFP+) and *Pdgfra* (membrane GFP-) transcripts (Figure 4C-D’).

**Figure 4.**
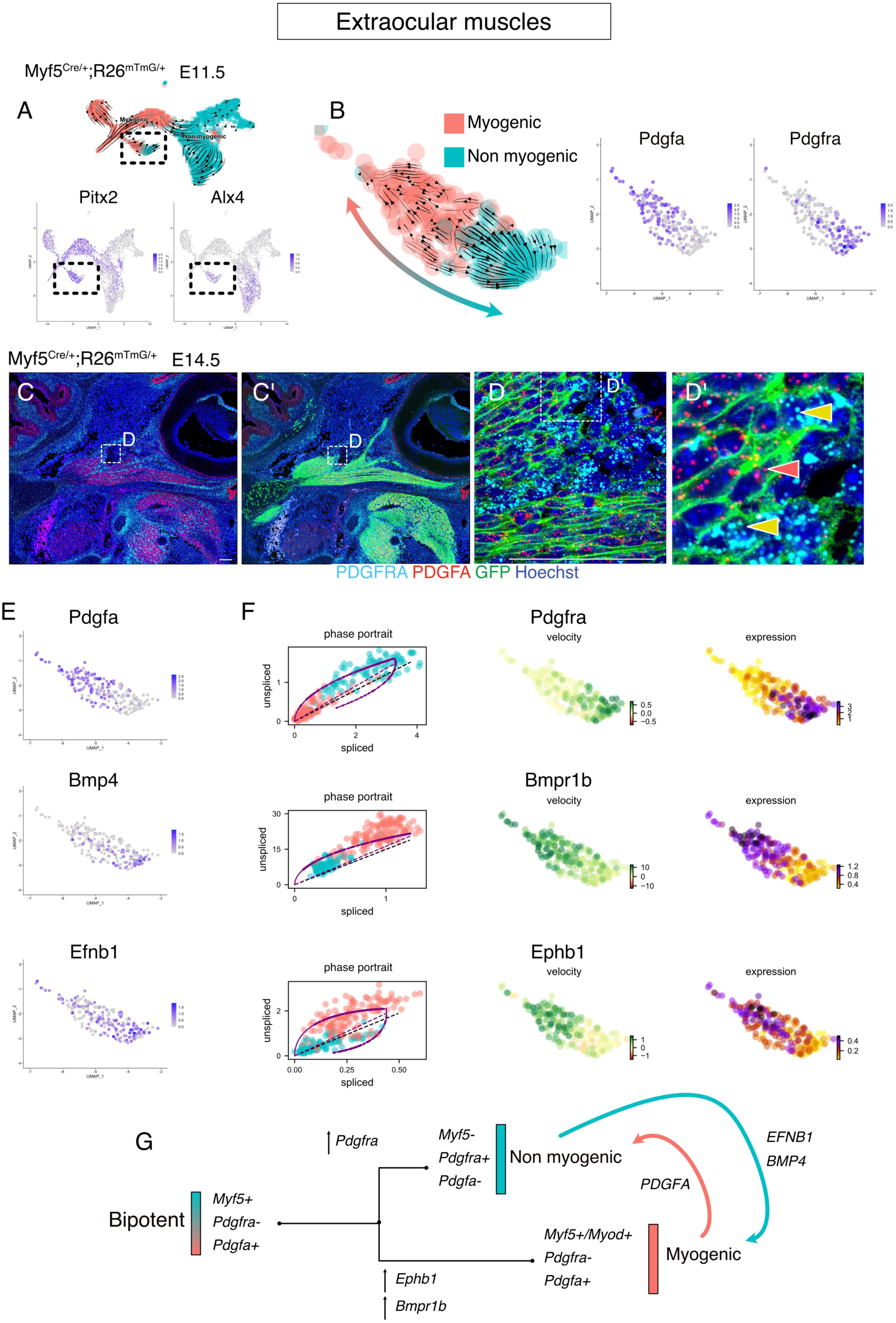
Maintenance of signalling cues between Myf5-derived myogenic and non-myogenic cells in EOM. A) UMAPs of *Myf5^Cre/+^; R26^mTmG/+^* E11.5 EOM subset and the respective RNA velocity trajectories (top). Expression plots of *Pitx2* and *Alx4*, marking the EOM cluster (bottom). B) Velocities within myogenic and non-myogenic clusters and expression plots of *Pdgfa* and *Pdgfra*. C-D’) RNAscope on *Myf5^Cre/+^; R26^mTmG/+^* E14.5 tissue sections with *Pdgfra* (cyan) and *Pdgfa* (red) probes. *Myf5*-derived cells are labelled by membrane GFP staining. D’) High magnification of D. Yellow arrowheads indicate *Myf5*-derived *Pdgfra*-expressing cells (non-myogenic). Red arrowheads indicate *Myf5*-derived *Pdgfa*-expressing cells (myogenic). E) Expression pattern of ligands *Pdgfa*, *Bmp4* and *Efnb1*. F) Phase portrait, gene velocity and expression plots of receptors *Pdgfra*, *Bmpr1b* and *Ephb1*. G) Model of myogenic and non-myogenic cell communication following bifurcation

Gene set enrichment analysis of EOM myogenic and non-myogenic driver genes revealed that transmembrane receptor protein kinase and SMAD activity were shared terms between the 2 clusters, indicating that specific complementary signalling networks could be actively maintained between these two populations (Figure S4A). Therefore, we examined the dynamic induction of tyrosine kinase ligands and receptors in the EOM. As observed in the scRNAseq dataset of anterior somites (Figure 2) and by RNAscope on tissue sections at the EOM level (Figure 4C-D’), *Pdgfra* expression was actively induced in non-myogenic *Myf5*-derived cells while *Pdgfa* expression was found in myogenic cells (Figure 4E-F). Notably, two additional tyrosine kinase receptors, namely *Bmpr1b* and *Ephb1,* were found to be among the top 100 driver genes of the myogenic EOM compartment, indicating that myogenic commitment is associated with upregulation of these receptors in the EOM (Figure 4F, Table1). Strikingly, two of their respective ligands *Bmp4* and *Efnb1*, were found to be specifically expressed in non-myogenic cells (Figure 4E). These results favor a model where complementary paracrine signalling networks operates between myogenic and non-myogenic *Myf5*-derived cells (Figure 4G), while their cellular juxtaposition is maintained through fetal stages.

### Obstructing myogenesis expands connective tissue formation from bipotent cells

The directional trajectories identified by RNA velocity in the EOM at E11.5 showed a strong bipolarity in fate with a higher velocity confidence index at each end of the myogenic and non-myogenic domains, and lower at their interface (Figure S4B). This suggests that the anticipated cell fate is bipotential before cell fate bifurcation. Conversely, cells that were located on either side of this central region were identified with greater confidence as committed either to a myogenic or a non-myogenic fate (Figure S4B). To identify the regulatory factors underlying this bipotency, we used SCENIC, a regulatory network inference algorithm (Aibar et al., 2017). This tool allows regrouping of sets of correlated genes into regulons (i.e. a transcription factor and its targets) based on binding motifs and co-expression. Use of this pipeline significantly reduced the number of variables from thousands of genes to a few hundred regulatory modules, while preserving the general aspect of the data, particularly the bipartite distribution of myogenic and non-myogenic cells (Figure 5A). The top regulons of this analysis revealed active transcription factors underlying myogenic and non-myogenic cell fates in the EOM at E11.5. Notably, *Myf5*, *Pitx1*, *Mef2a* and *Six1*, transcription factors known to be implicated in myogenic development (Buckingham and Rigby, 2014), appeared among the top regulons in myogenic cells whereas *Fli1*, *Ebf1*, Ets1, *Foxc1*, *Meis1* and *Six2*, genes known for their involvement in adipogenic, vascular, mesenchymal and tendon development (Jimenez et al., 2006; López-Delgado et al., 2020; Noizet et al., 2016; Truong and Ben-David, 2000; Whitesell et al., 2019; Yamamoto-Shiraishi and Kuroiwa, 2013), constituted some of the highly active non-myogenic transcription factors (Figure 5B). Given that *Myf5* appeared as a top regulatory factor of the myogenic program, we interrogated the fate of *Myf5*-expressing progenitors in a *Myf5^nlacZ/nlacZ^* null genetic background. Interestingly, some β-gal+ cells were found in the cartilage primordium (Sox9+) of the EOM in the heterozygous control at E12.5 indicating that cells with recent *Myf5* activity diverged to a non-myogenic fate (Figure 5C-C’). As previously reported, the extraocular muscles are absent in this mutant (Figure 5D, asterisk) (Sambasivan et al., 2009). Notably, disruption of *Myf5* activity led to a 3-fold increase in the proportion of non-myogenic *Myf5*-derived cells in this region (Figure 5E). In contrast, no double-positive cells were found in the masseter, a muscle fully embedded in neural crest, even in the absence of *Myf5* (Figure 5E). Robust *Myf5* expression is thus necessary to maintain a balance between myogenic and non-myogenic cell fates of Myf5+ bipotent progenitors only in neural crest-depleted regions. Conversely, virtually no Pdgfra+ cells were found to be derived from *Myod* expressing cells in most muscles of *Myod^iCre^;R26^TdTomato/+^;Pdgfra^H2BGFP/+^* fetuses at E14.5, and only rare double positive cells were found in the EOM (Figure S5). This observation indicates that bipotency is associated with *Myf5*, and that subsequent activation of *Myod* within this lineage locks cell fate into the myogenic program thereby suppressing their connective tissue potential (Figure 5F).

**Figure 5.**
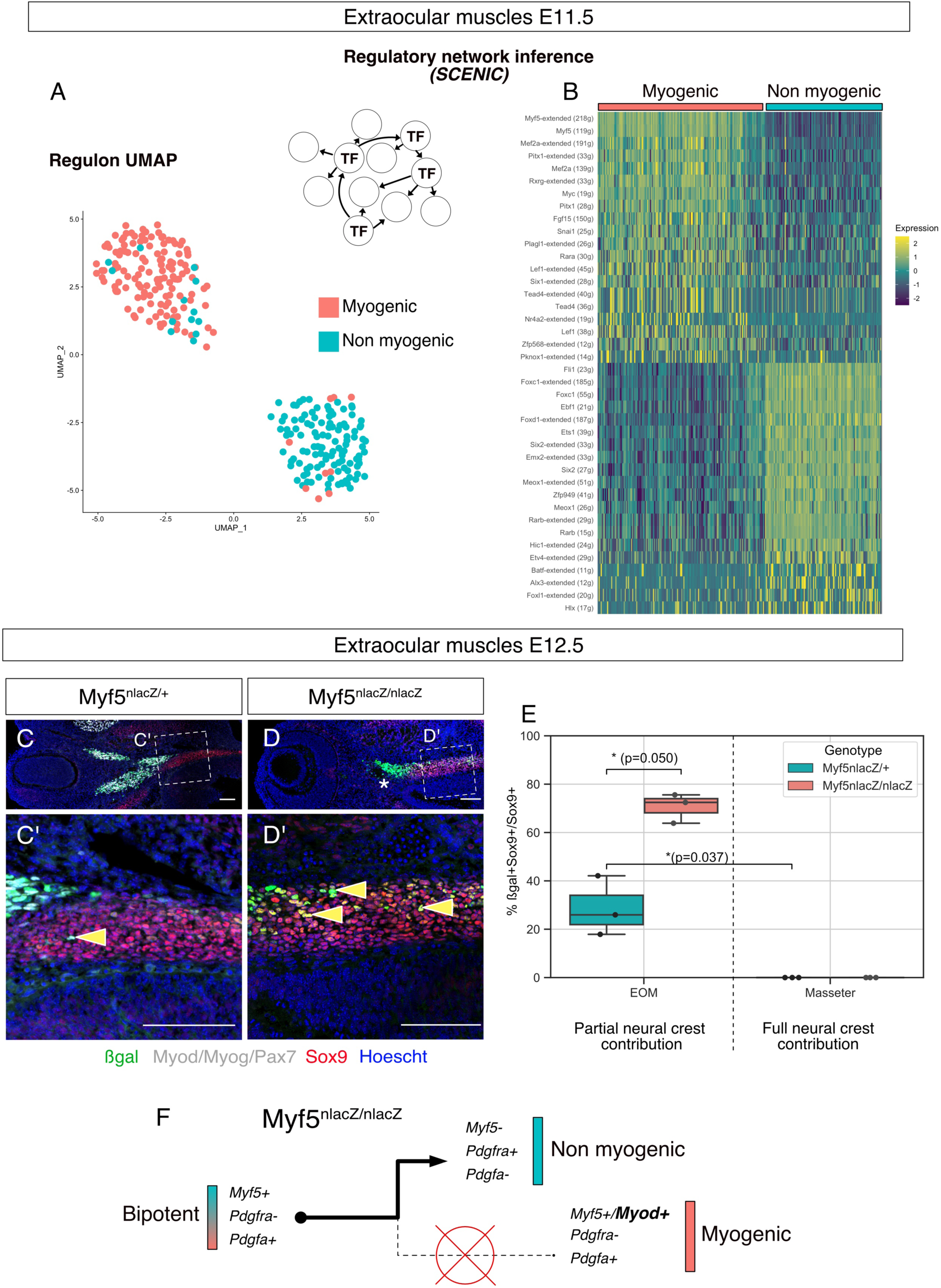
Disruption of *Myf5* increases the connective tissue output from bipotent cells. A) UMAP of *Myf5^Cre/+^; R26^mTmG/+^* E11.5 EOM based on SCENIC Regulon activity (Area Under Curve score). B) Heatmap of top regulons (transcription factor and associated targets). The suffix “_extended” indicates that the regulon includes motifs that have been linked to the TF by lower confidence annotations, for instance, inferred by motif similarity. Number in brackets indicates number of genes comprising the regulon. C-D) Transverse sections of *Myf5^nlacZ/+^* (C-C’), and *Myf5^nlacZ/nlacZ^* (D-D’) in the EOM region at E12.5 immunostained for β-gal (green), Sox9 (red) and Myod/Myog/Pax7 (grey). Yellow arrowheads indicate β-gal/Sox9 double positive cells and show an expansion these cells in the mutant. Asterisk highlights the lack of myogenic progenitors in the EOM region of the mutant embryo, indicated by the absence of Myod/Myog/Pax7 staining. E) Quantification of proportion of β-gal+;Sox9+ double positive cells in the total Sox9+ population of the EOM and Masseter muscles. Each dot is a different sample, the center line of the boxplot is the median value. (n=3 embryos, p-values were calculated using a two-sided Mann-Whitney U test). F) Model of lineage progression from bipotent cells in a *Myf5* null background.

### Myf5-derived contribution to connective tissues is sustained through muscle initiation

Although we identified *Myf5*-derived non-myogenic cells in various regions of the embryo, it was not clear if this population was self-sustaining, or continuously generated throughout development. To address this issue, we performed 2 more scRNAseq experiments at E12.5 and E14.5, using contemporary *Myf5* labelling (*Myf5^GFP-P/+^*; Figures 6, S1B, G-J, S3A, D-E). In accordance with the earlier datasets, cells that appeared to belong to muscle anlagen of EOM, somites and caudal arches progressed towards a non-myogenic state (Figure 6A-C’). To assess the identity of these cells, we performed a gene set enrichment network analysis combining the differentially expressed genes of non-myogenic clusters of all stages. We found that all stages contributed equally to “GO Molecular Function” and “Reactome pathways” terms in spite of their relatively diverse gene expression signatures (Figure S6). This finding suggests that these non-myogenic cells are relatively homogeneous in gene signatures throughout cranial muscles when they emerge from common bipotent progenitors. Highly significant terms hinted at a myogenic-supporting role, providing muscle progenitors with extracellular matrix components, and contributing to neuronal guidance (Figure 6E). Among these terms, presence of Pdgf signalling and receptor kinase activity indicated, once again, that the interactions found in the EOM could occur also at later stages in various craniofacial muscles that are deprived of neural crest derived connective tissue.

**Figure 6.**
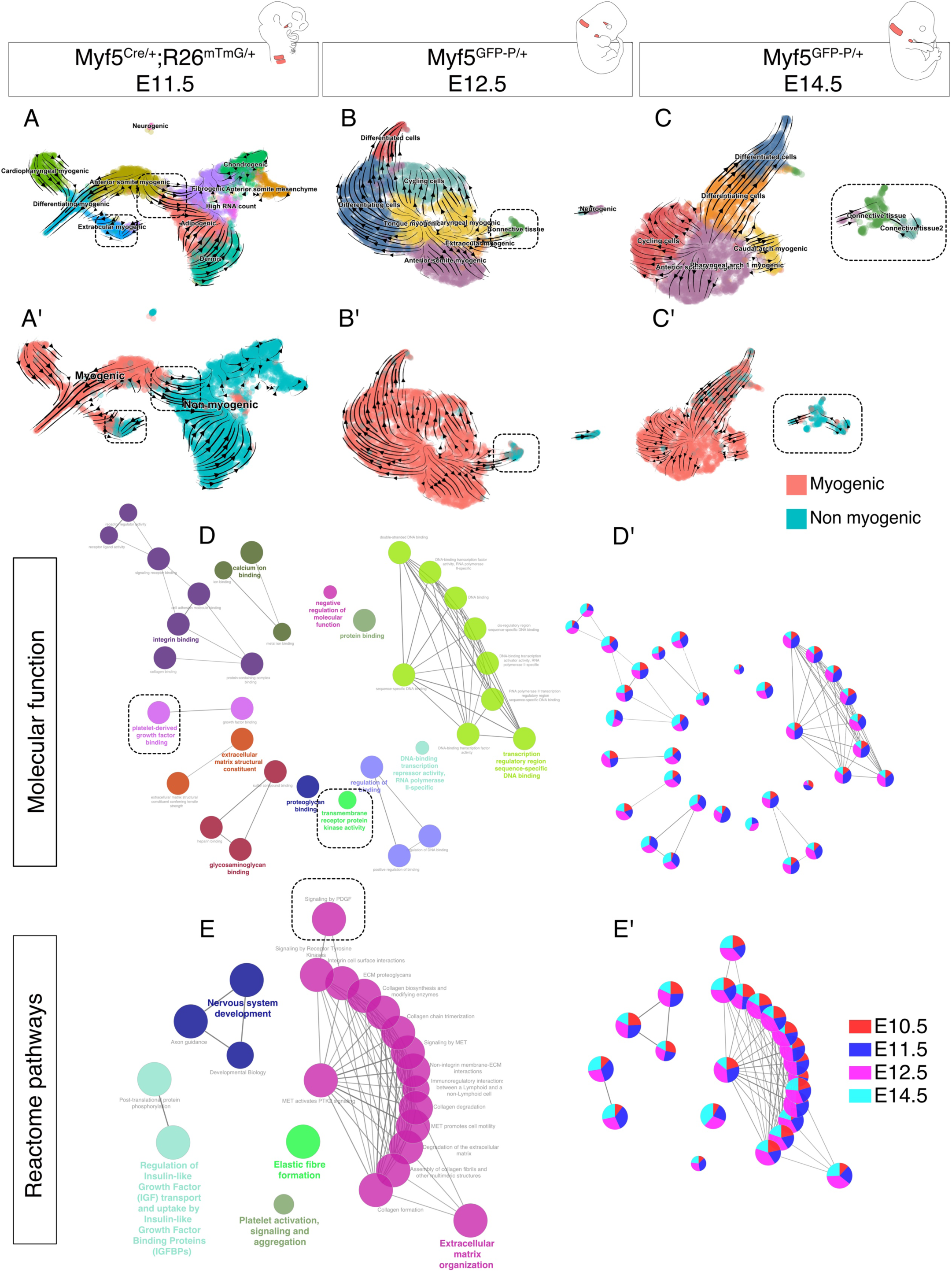
Myf5-derived non-myogenic cells are continuously generated up to fetal stages. A-C’) RNA velocity plots of *Myf5^Cre/+^; R26^mTmG/+^* E11.5, *Myf5^GFP-P/+^* E12.5 and *Myf5^GFP-P/+^* E14.5 datasets displaying cell-type annotation (A-C) and myogenic and non-myogenic clustering (A’-C’). D-E) Gene ontology network of GO Molecular Function and Reactome pathway performed on combined top 100 markers. (D’-E’) Relative contribution of each stage to term node.

### A novel regulatory network underlies the non-myogenic cell fate

Myf5+ bipotent progenitors were observed at multiple stages and anatomical locations, and they yielded a relatively homogeneous population expressing common markers associated with extracellular matrix components, cell adhesion molecules, and tyrosine kinase signalling. To assess whether the regulatory mechanisms guiding this transition are distinct in different locations in the head, we set out to explore the common molecular switches underlying cell fate decisions. To do so, we developed a pipeline where we combined the list of driver genes at the start of the non-myogenic trajectory (Table 1) with the most active regulons in the non-myogenic region (Methods, code in open access). This resulted in a network consisting of the most active transcription factors and the most transcriptionally dynamic genes found at the non-myogenic branchpoint. We performed this operation for each dataset independently and displayed them as individual networks (Figure 7A-D). Finally, we overlapped the list of these “driver regulators” to identify the common transcription factors guiding the non-myogenic cell fate decision (Figure 7E). Notably, *Foxp2*, *Hmga2*, *Meis1*, *Meox2* and *Tcf7l2* were identified in all 4 datasets as key driver regulators, and thus are likely to play significant role in the non-myogenic transition (Figure 7E, Table 2).

**Figure 7.**
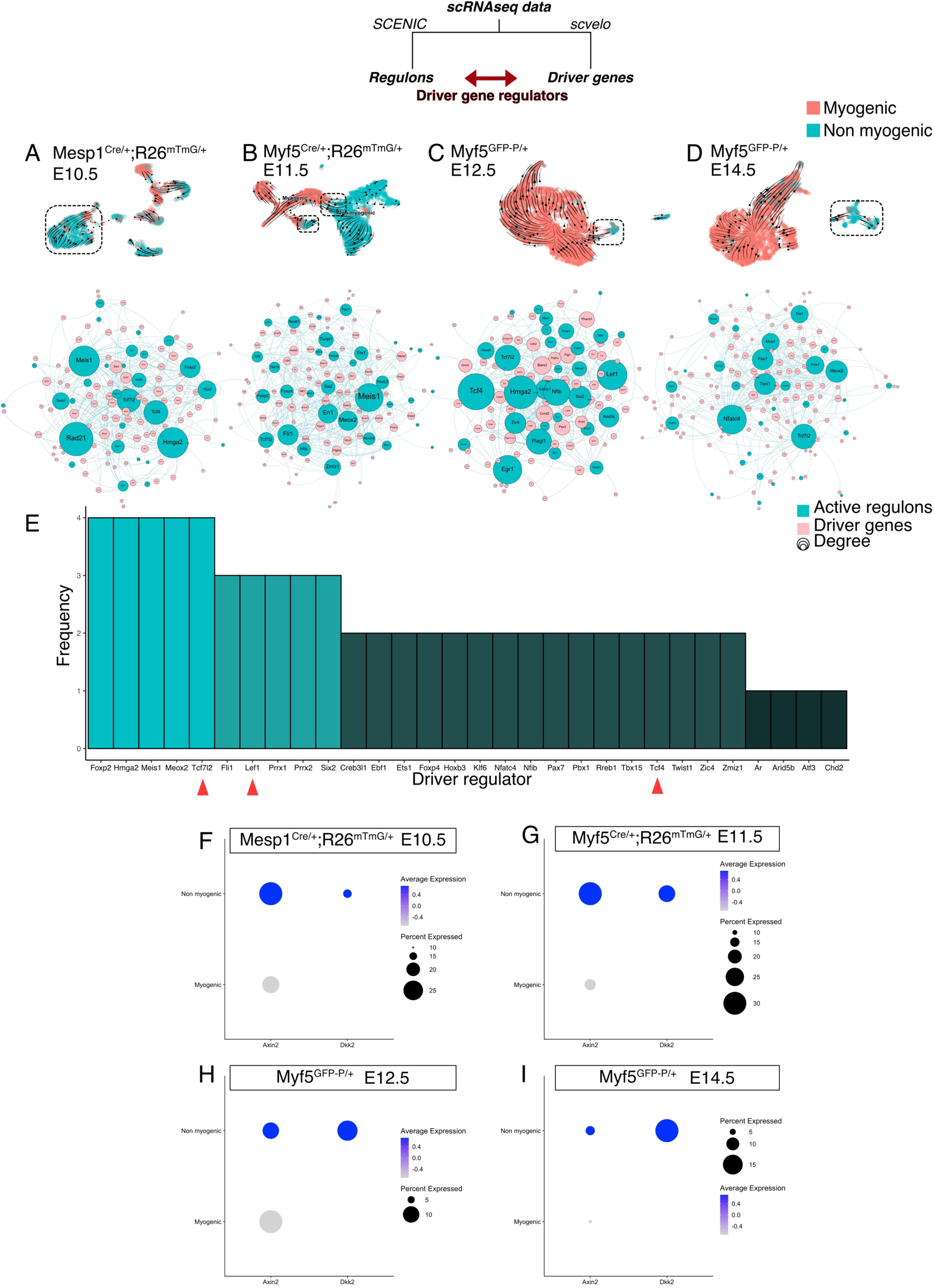
A shared program including Wnt/β-cat activity supports non-myogenic fate transition at various stages and anatomical locations. A-D) Velocity UMAP highlighting the non-myogenic cluster at each stage (dotted box), from which the underlying network was inferred. Driver genes and regulatory networks (regulons) were produced for each stage independently, and a stage-specific network of active transcription factor and associated driver gene targets was built. Size of nodes corresponds to the number of edges (connections) they have, i.e. the number of driver genes the transcription factor regulates. E) Histogram displaying frequency of appearance of most predominant transcription factors as driver regulators (4= present in all 4 datasets as driver regulon, 1= present in a single dataset). Red arrowheads highlight members of the Wnt/β-cat pathway Tcf4, Tcf7l2 and Lef1. F-I) Dotplot of the expression levels and percent of Axin2 and Dkk2 in the myogenic and the non-myogenic portions of all 4 datasets.

**Table 2:** Driver regulators of non-myogenic fate in each dataset. (+): Present, (-): Absent.

|  | E10.5 | E11.5 | E12.5 | E14.5 |
| --- | --- | --- | --- | --- |
| Foxp2 | (+) | (+) | (+) | (+) |
| Hmga2 | (+) | (+) | (+) | (+) |
| Meis1 | (+) | (+) | (+) | (+) |
| Meox2 | (+) | (+) | (+) | (+) |
| Tcf7l2 | (+) | (+) | (+) | (+) |
| Fli1 | (+) | (+) | (+) | (-) |
| Lef1 | (-) | (+) | (+) | (+) |
| Prrx1 | (+) | (+) | (-) | (+) |
| Prrx2 | (-) | (+) | (+) | (+) |
| Six2 | (+) | (+) | (+) | (-) |
| Creb3l1 | (-) | (+) | (-) | (+) |
| Ebf1 | (+) | (-) | (+) | (-) |
| Ets1 | (-) | (+) | (-) | (+) |
| Foxp4 | (+) | (+) | (-) | (-) |
| Hoxb3 | (-) | (+) | (+) | (-) |
| Klf6 | (-) | (+) | (-) | (+) |
| Nfatc4 | (-) | (+) | (-) | (+) |
| Nfib | (-) | (+) | (+) | (-) |
| Pax7 | (-) | (-) | (+) | (+) |
| Pbx1 | (-) | (+) | (-) | (+) |
| Rreb1 | (-) | (-) | (+) | (+) |
| Tbx15 | (+) | (+) | (-) | (-) |
| Tcf4 | (+) | (-) | (+) | (-) |
| Twist1 | (+) | (+) | (-) | (-) |
| Zic4 | (+) | (-) | (+) | (-) |
| Zmiz1 | (-) | (+) | (+) | (-) |
| Ar | (-) | (-) | (-) | (+) |
| Arid5b | (-) | (-) | (+) | (-) |
| Atf3 | (-) | (-) | (-) | (+) |
| Chd2 | (+) | (-) | (-) | (-) |

Interestingly, Tcfs and Lef1 were among these top common regulators and they form a complex effector for the canonical Wnt pathway. Previous work showed that during cranial myogenesis, neural crest cells release inhibitors of the Wnt pathway to promote myogenesis (Tzahor et al., 2003). It is thus tempting to speculate that in the absence of neural crest, mesoderm-derived bipotent progenitors can give rise to connective tissue by maintaining canonical Wnt activity. To assess this hypothesis, we examined the expression of Axin2, a common readout for Wnt/β-cat activity (Babb et al., 2017; Moosdijk et al., 2020). Interestingly, Axin2 levels were elevated in the non-myogenic portion of all the different datasets (Figure 7F-I). Additionally, Dkk2, which has been described as an activator of Wnt/β-cat pathway in the neural crest (Devotta et al., 2018), was also found to be elevated, indicative of a putative positive-feedback loop mechanism supporting the maintenance of this population.

## DISCUSSION

Distinct fates can emerge through the specification of individual cell types, or through direct lineage ancestry from bipotent or multipotent cells. Here we addressed this issue in the context of the emergence of myogenic and associated connective tissue cells during the formation of craniofacial muscles. By combining state-the-art analytical methods, we identified the transcriptional dynamics, the intercellular communication networks, and the regulators controlling the balance between two complementary cell fates. Specifically, our work provides evidence for a novel mesoderm-derived bipotent cell population that gives rise to muscle and associated connective tissue cells spatiotemporally, and only in regions deprived of neural crest cells (Figure S7).

Brown adipocytes, neurons, pericytes and rib cartilage have been reported to express *Myf5* in ancestral cells (Daubas et al., 2000; Haldar et al., 2008; Sebo et al., 2018; Stuelsatz et al., 2014). Interestingly, when *Myf5* expression is disrupted, cells can acquire non-myogenic fates and contribute to connective tissue (this study), cartilage, and dermis (Tajbakhsh et al., 1996). These studies suggest that *Myf5*- expression alone is not sufficient to promote robust myogenic fate in multiple regions of developing embryos. Consistent with these observations, Myod+ cells do not contribute to rib cartilage (Wood et al., 2020) and only give rise to very few connective tissue cells in the periocular region (this study). These findings are also consistent with the higher chromatin-remodelling capacity of Myod compared to Myf5, and its role as defining the committed myogenic cell state (Conerly et al., 2016; Tapscott, 2005). In contrast to a previous study (Stuelsatz et al., 2014), we found no neural-crest derived cells expressing *Myf5* during EOM tissuegenesis at E13.5 (using *Wnt1^Cre/+^;R26^mTmG/+^;Myf5^nlacZ/+^*). We note that *Myf5*- expressing cells contribute to non-myogenic cells from early embryonic stages (E10.5) and continue to do so in the fetus, indicating that these bipotent cells persist well after muscles are established.

Here, we also identifed a core set of transcription factors specifically active in the non-myogenic cells across all datasets. We propose that these genes guide bipotent cells to a non-myogenic fate and thus confer mesenchymal properties to non-committed progenitors. Recent studies have identified anatomically distinct fibroblastic populations using single-cell transcriptomics, yet unique markers could not be identified (Muhl et al., 2020; Sacchetti et al., 2016), making characterisation of cell subtypes challenging. Tcf4/Tcf7l2 was identified as a master regulator of fibroblastic fate during muscle-associated connective tissue development although it is also expressed in myogenic progenitors at lower levels (Kardon et al., 2003; Mathew et al., 2011; Sefton and Kardon, 2019). We also report that this gene is one of the main regulators of connective tissue fate. Other transcription factors have been linked to skin fibroblast fates including *Tcf4*, *Six2*, *Meox2*, *Egr2* and *Foxs1*, and their repression favours a myofibroblastic potential (Noizet et al., 2016). *Six2* and *Meox2* were also identified in our analysis, which raises the question of the shared genetic programs between myofibroblastic cells and fibroblastic cells derived from progenitors primed for myogenesis during development.

Interestingly, *Prrx1*, a marker for lateral plate mesoderm (Durland et al., 2008), was differentially expressed in the connective tissue population at various stages. Although lateral plate mesoderm is identifiable in the trunk, its anterior boundaries in the head are unclear (Prummel et al., 2020). More detailed analyses of *Prrx1*, *Isl1* and *Myf5* lineages need to be carried out to delineate the specific boundaries of each progenitor contribution to cranial connective tissue.

Tyrosine kinase receptors have been implicated in a number of developmental programs for both muscle and associated connective tissues (Arnold et al., 2020; Knight and Kothary, 2011; Olson and Soriano, 2009; Tallquist et al., 2000; Tzahor et al., 2003; Vinagre et al., 2010). For example, the differentiation of fetal myoblasts is inhibited by growth factors Tgfβ and Bmp4 (Cossu et al., 2000). Epha7 signalling is expressed in embryonic and adult myocytes and promotes their differentiation (Arnold et al., 2020). Significantly, we noticed a striking and lasting complementary expression of *Pdgfa* and *Pdgfra* throughout embryonic stages, in the myogenic and non-myogenic progenitors respectively. Pdgf ligands emanating from hypaxial myogenic cells under the control of *Myf5* were shown to be necessary from rib cartilage development (Tallquist et al., 2000; Vinagre et al., 2010). Additionally, Pdgfra promotes expansion of fibroblasts during fibrosis (Olson and Soriano, 2009). Interestingly, we found that *Pdgfa* expression was reduced in cells expressing high levels of *Myog* at the fetal stage (data not shown). Therefore, *Myf5*-derived myogenic progenitor cells might guide non-myogenic *Myf5*-derived expansion, which in turn provides ligands and extracellular matrix components to favour myogenic development and patterning. Moreover, unlike trunk myogenesis, cranial muscle development relies on the expression of Wnt and Bmp inhibitors from surrounding tissues (Tzahor et al., 2003). Interestingly, we showed that the *Myf5*-derived non-myogenic cells express Bmp4, Dkk2, and Axin2. Additionally, we showed that the Wnt effector complex Tcf/Lef is active in these cells. It is thus likely that these cells maintain their non-myogenic fate by promoting Bmp production and Wnt activity cell-autonomously. Further studies could provide further insights into the evolutionary ancestry of this bipotency by studying other model organisms devoid of neural crest.

## MATERIALS & METHODS

### scRNAseq data generation

For E10.5 to E12.5 embryos, the cranial region above the forelimb was dissected in ice-cold 3% FBS in PBS and mechanically dissociated with forceps and pipetting. The same procedure was applied at E14.5 but the dissection was refined to the pharyngeal and laryngeal regions. Tissues were then digested in TrypLE (ThermoFisher Cat #: 12604013) during 3 rounds of 5-min incubation (37°C, 1400 RPM), interspersed with gentle pipetting to further dissociate the tissue. Cells were resuspended in FBS 3%, filtered, and incubated with Calcein Blue (eBioscience, Cat #: 65-0855-39) and Propidium Iodide (ThermoFisher Cat #: P1304MP) to check for viability. Viable cells were sorted on BD FACSAria™ III and manually counted using a hemocytometer. RNA integrity was assessed with Agilent Bioanalyzer 2100 to validate the isolation protocol prior to scRNAseq (RIN>8 was considered acceptable). 4000 to 13000 cells were loaded onto 10X Genomics Chromium microfluidic chip and cDNA libraries were generated following manufacturer’s protocol. Concentrations and fragment sizes were measured using Agilent Bioanalyzer and Invitrogen Qubit. cDNA libraries were sequenced using NextSeq 500 and High Output v2.5 (75 cycles) kits. Genome mapping and count matrix generation were done following 10X Genomics Cell Ranger pipeline.

### RNA velocity and driver genes

RNA velocity analyses were performed using scvelo (Bergen et al., 2020) in python. This tool allows inferring velocity flow and driver genes using scRNAseq data, with major improvements from previous methods (Manno et al., 2018). First, unspliced and spliced transcript matrices were generated using velocyto (Manno et al., 2018) command line function, which outputs unspliced, spliced, and ambiguous matrices as a single loom file. These files were combined with filtered Seurat objects to yield objects with unspliced and spliced matrices, as well as Seurat-generated annotations and cell-embeddings (UMAP, tSNE, PCA). These datasets were then processed following scvelo online guide and documentation. Velocity was calculated based on the dynamical model (using *scv.tl.recover_dynamics(adata)*, and *scv.tl.velocity(adata, mode=’dynamical’)*) and when outliers were detected, differential kinetics based on top driver genes were calculated and added to the model (using *scv.tl.velocity(adata, diff_kinetics=True)*). Specific driver genes were identified by determining the top likelihood genes in the selected cluster. The lists of top 100 drivers for each stage are given in Table1.

### Seurat preprocessing

scRNAseq datasets were preprocessed using Seurat in R (https://satijalab.org/seurat/) (Butler et al., 2018). Cells with more than 20% of mitochondrial gene fraction were discarded. The number of genes expressed averaged to 4000 in all 4 datasets. Dimension reduction and UMAP generation were performed following Seurat workflow. Doublets were inferred using DoubletFinder v3 (McGinnis et al., 2019). Cell cycle genes, mitochondrial fraction, number of genes, number of UMI were regressed in all datasets following Seurat dedicated vignette. We noticed that cell cycle regression, although clarifying anatomical diversity, seemed to induce low and high UMI clustering (Figure S1E-F). For the E10.5 and E11.5 datasets, 2 replicates were generated from littermates and merged after confirming their similitude. For subsequent datasets (E12.5 and E14.5), no replicates were used. Annotation and subsetting were also performed in Seurat. “Myogenic” and “Non-myogenic” annotations were based on *Pdgfa* and *Pdgfra* expression and myogenic genes *Myf5*, *Myod*, and *Myog*. Cells not expressing *Pdgfa* were annotated as “non-myogenic” unless they express myogenic genes. Cells expressing *Pdgfa* were annotated as “myogenic”. We noticed that at later stages, *Pdgfa* expression decreases in Myog+ cells. Thus, driver genes of connective tissue at E12.5 and E14.5 were determined using cluster annotations obtained from Leiden-based clustering.

### Gene regulatory network inference

Gene regulatory networks were inferred using SCENIC (R implementation) (Aibar et al., 2017) and pySCENIC (python implementation) (Sande et al., 2020). This algorithm allows regrouping of sets of correlated genes into regulons (i.e. a transcription factor and its targets) based on motif binding and co-expression. UMAP and heatmap were generated using regulon AUC matrix (Area Under Curve) which refers to the activity level of each regulon in each cell.

### Driver regulons

Results from SCENIC and scvelo were combined to identify regulons that could be responsible for the transcriptomic induction of driver genes. Similar to the steps mentioned above, SCENIC lists of regulons were used to infer connections between transcription factors and driver gene. Networks were generated as explained above, and annotated with “Active regulon” or “driver gene”. The lists of individual driver regulons of each dataset were then combined and the most recurring driver regulons were identified. The code is available at this address: https://github.com/TajbakhshLab/DriverRegulators

### Gene set enrichment analysis

Gene set enrichment analyses were performed on either the top markers (obtained from Seurat function FindAllMarkers) or from driver genes (obtained from scvelo), using Cluego (Bindea et al., 2009). “GO Molecular Pathway”, “GO Biological Process” and “Reactome pathways” were used independently to identify common and unique pathways involved in each dataset. In all analyses, an enrichment/depletion two-sided hypergeometric test was performed and p-values were corrected using the Bonferroni step down method.

### Mouse strains

Animals were handled according to European Community guidelines and the ethics committee of the Institut Pasteur (CETEA) approved protocols. The following strains were previously described: *Myf5^Cre^* (Haldar et al., 2008), *Mesp1^Cre^* (Saga et al., 1999), *Tg:Wnt1Cre* (Danielian et al., 1998), *R26^TdTom^* (Ai9;(Madisen et al., 2009)), *R26^mTmG^*(Muzumdar et al., 2007), *Myf5^nlacZ^* (Tajbakhsh et al., 1996), *Pdgfra^H2BGFP^* (Hamilton et al., 2003) and *Myf5^GFP-P^* (Kassar-Duchossoy et al., 2004). To generate *Myf5^Cre/+^;R26^TdTomato/+^;Pdgfra^H2BGFP/+^*embryos, *Myf5^Cre/+^* females were crossed with *Pdgfra^H2BGFP/+^;R26^TdTomato/TdTomato^* males. Mice were kept on a mixed genetic background C57BL/6JRj and DBA/2JRj (B6D2F1, Janvier Labs). Mouse embryos and fetuses were collected between embryonic day (E) E10.5 and E14.5, with noon on the day of the vaginal plug considered as E0.5.

### Immunofluorescence

Collected embryonic and adult tissues were fixed 2.5h in 4% paraformaldehyde (Electron Microscopy Sciences, Cat #:15710) in PBS with 0,2-0,5% Triton X-100 (according to their stage) at 4°C and washed overnight at 4°C in PBS. In preparation for cryosectioning, embryos were equilibrated in 30% sucrose in PBS overnight at 4°C and embedded in OCT. Cryosections (16-20µm) were left to dry at RT for 30 min and washed in PBS. The primary antibodies used in this study are chicken polyclonal anti-β-gal (Abcam, Cat #: ab9361, dilution 1:1000), mouse monoclonal IgG1, mouse monoclonal IgG1 anti-Myod (BD Biosciences, Cat# 554130, dilution 1:100), mouse monoclonal IgG1 anti-Pax7 (DSHB, Cat. #: AB_528428, dilution 1:20), rabbit anti-mouse Sox9 (Millipore, Cat. #: AB5535, dilution 1/2000), rabbit polyclonal anti-Tomato (Clontech Cat. #: 632496, dilution 1:400) and chicken polyclonal anti-GFP (Abcam Cat. #: 13970, dilution 1:1000). Images were acquired using Zeiss LSM780 or LSM700 confocal microscopes and processed using ZEN software (Carl Zeiss).

### RNAscope in situ hybridization

Embryos for in situ hybridization were fixed overnight in 4% PFA. Embryos were equilibrated in 30% sucrose in PBS and sectioned as described for immunofluorescence. RNAscope probes Mm-Pdgfa (411361) and Mm-Pdgfra (480661-C2) were purchased from Advanced Cell Diagnostics, Inc. In situ hybridization was performed using the RNAscope Multiplex Fluorescent Reagent Kit V2 as described previously (Comai et al., 2019).

### Data Availability

The data that support the findings of this study are available from the corresponding author, S.T, upon request. The code that was used to generate the driver regulators is available at this address: https://github.com/TajbakhshLab/DriverRegulators

## Acknowledgements

We acknowledge funding support from the Institut Pasteur, Association Française contre le Myopathies, Agence Nationale de la Recherche (Laboratoire d’Excellence Revive, Investissement d’Avenir; ANR-10-LABX-73) and MyoHead, Association Française contre les Myopathies (Grant #20510), Fondation pour la Recherche Médicale (Grant # FDT201904008277), and the Centre National de la Recherche Scientifique. We gratefully acknowledge the UtechS Photonic BioImaging, C2RT, Institut Pasteur, supported by the French National Research Agency (France BioImaging; ANR-10–INSB–04; Investments for the Future).

## SUPPLEMENTAL INFORMATION

**Supplemental Figure S1.**
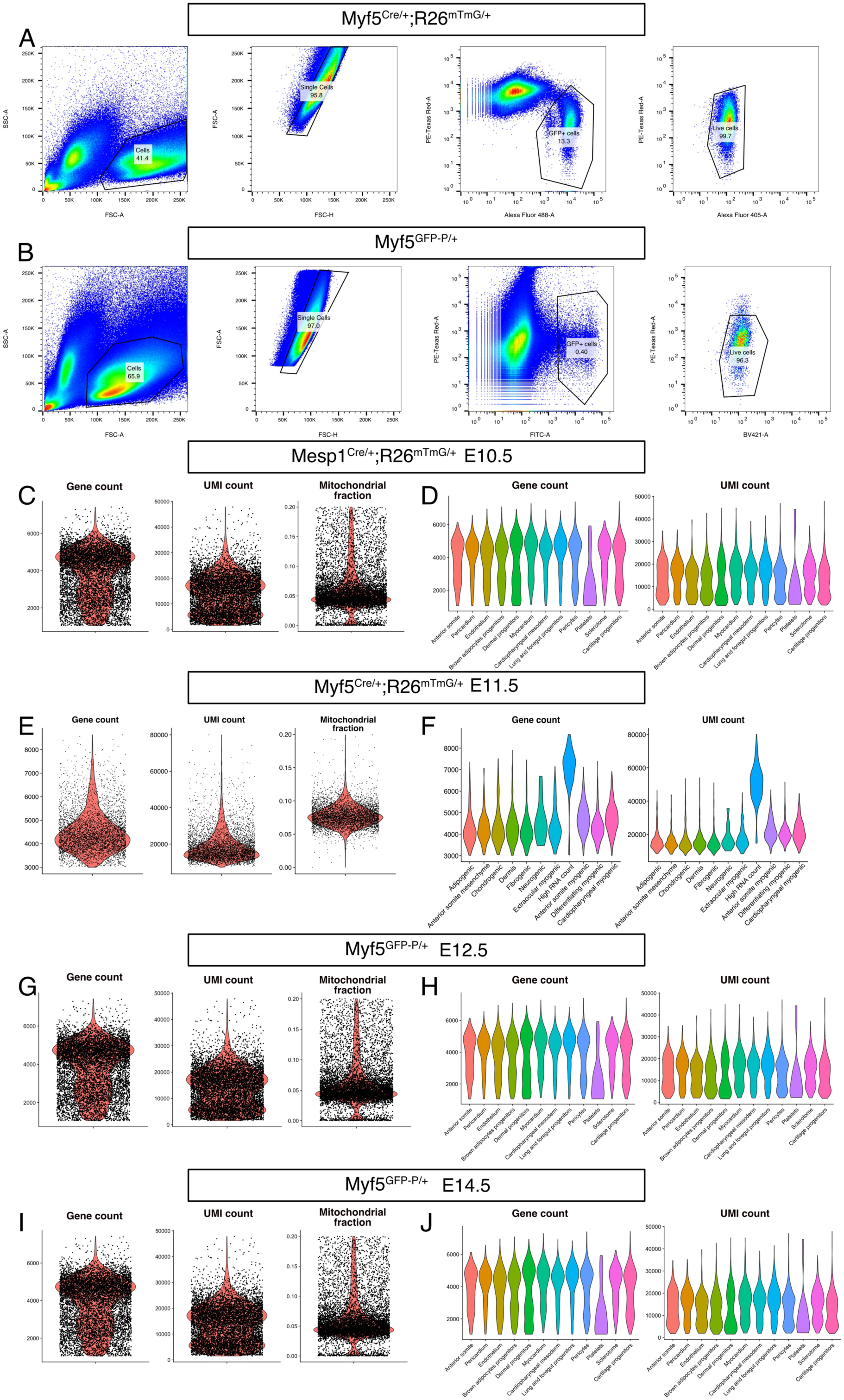
Seurat library preprocessing metrics. A-B) Gating strategy used to isolate by FACS *Myf5^Cre/+^; R26^mTmG/+^*(A) and *Myf5^GFP-P/+^* cells (B). To isolate *Mesp1^Cre/+^; R26^mTmG/+^* cells, the same strategy as in (A) was used. The Alexa Fluor 488 and FITC channels were used interchangeably to identify GFP+ cells. The BV421 and Alexa Fluor 405 were used interchangeably to identify the Calcein Blue+ live cells. The PE-Texas Red channel was used to discard mTomato+ cells and Propidium Iodide + cells. The percentage of cells captured by each gate is displayed on each plot. C, E, G, I) Violin plots of gene count, UMI count and mitochondrial fraction for each dataset. D, F, H, J) Gene count and UMI count per cell type for each dataset. A “High count” cluster was found in the E11.5 dataset, and was removed in downstream analysis.

**Supplemental Figure S2.**
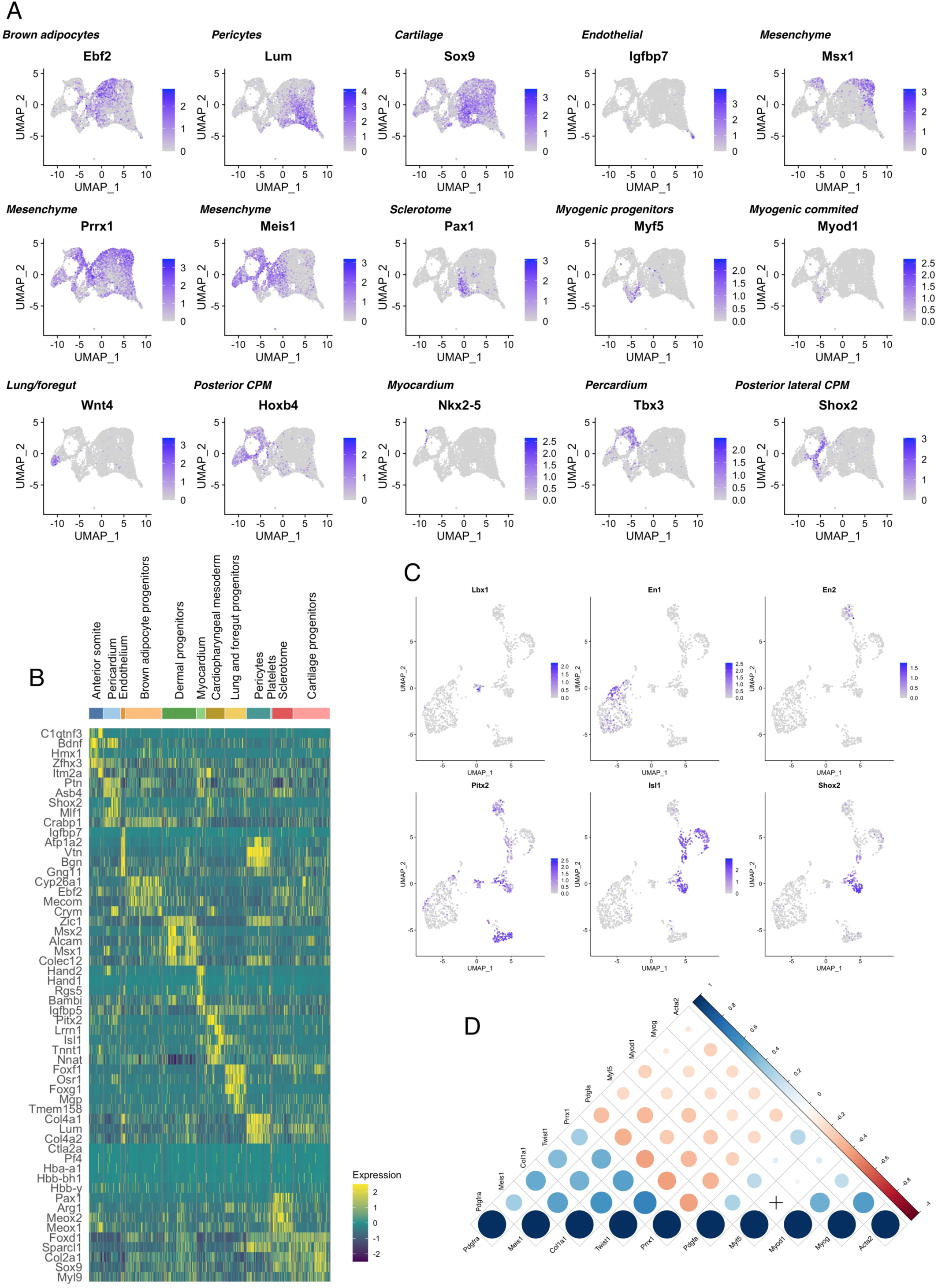
Myogenic and non-myogenic markers define anterior mesodermal tissues. A) *Mesp1^Cre/+^;R26^mTmG/+^* E10.5 UMAP expression plots of markers of various mesodermal lineages. B) Heatmap of top 5 markers of each cluster of *Mesp1^Cre/+^;R26^mTmG/+^* E10.5. C) UMAP expression plot of the *Mesp1^Cre/+^;R26^mTmG/+^* E10.5 subset. *En2*: marker of pharyngeal arch 1 (Knight et al., 2008), *En1*: marker of epaxial somitic progenitors (Cheng et al., 2004), *Lbx1*: marker for tongue progenitors (Gross et al., 2000), *Isl1*: marker of cardiopharyngeal mesoderm of pharyngeal arch 2-6 (Comai et al., 2019), *Shox2*: marker of caudal cardiopharyngeal mesoderm (Wang et al., 2020), *Pitx2*: marker of the extraocular region (Zacharias et al., 2010). D) Pearson correlation plot of myogenic (*Pdgfa, Myf5, Myod1, Myog, Acta2*) and non-myogenic (*Pdgfra, Prrx1, Meis1, Twist1, Osr1, Col1a1*) genes. The size of the dots is inversely proportional to their p-value. A cross indicates a p-value higher than 0.05. The color of the dots indicates the strength of the a positive (blue) or negative (red) correlation.

**Supplemental Figure S3.**
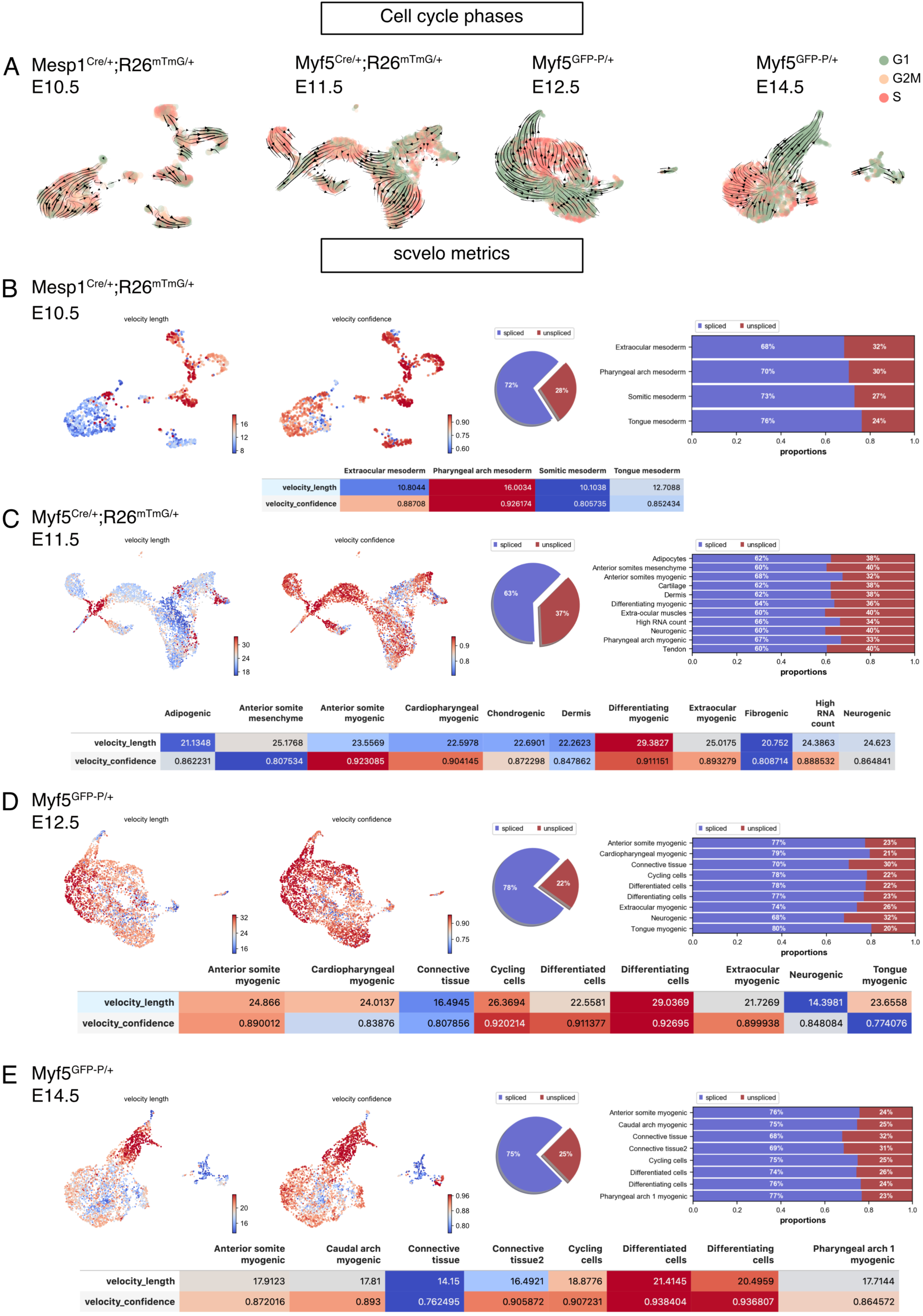
Cell cycle phases and scvelo metrics. A) UMAP of each dataset with overlaid velocity and cell cycle phase. B-E) Quality control metrics of scvelo, including velocity length, velocity confidence and spliced/unspliced abundance per dataset and cell type.

**Supplemental Figure S4.**
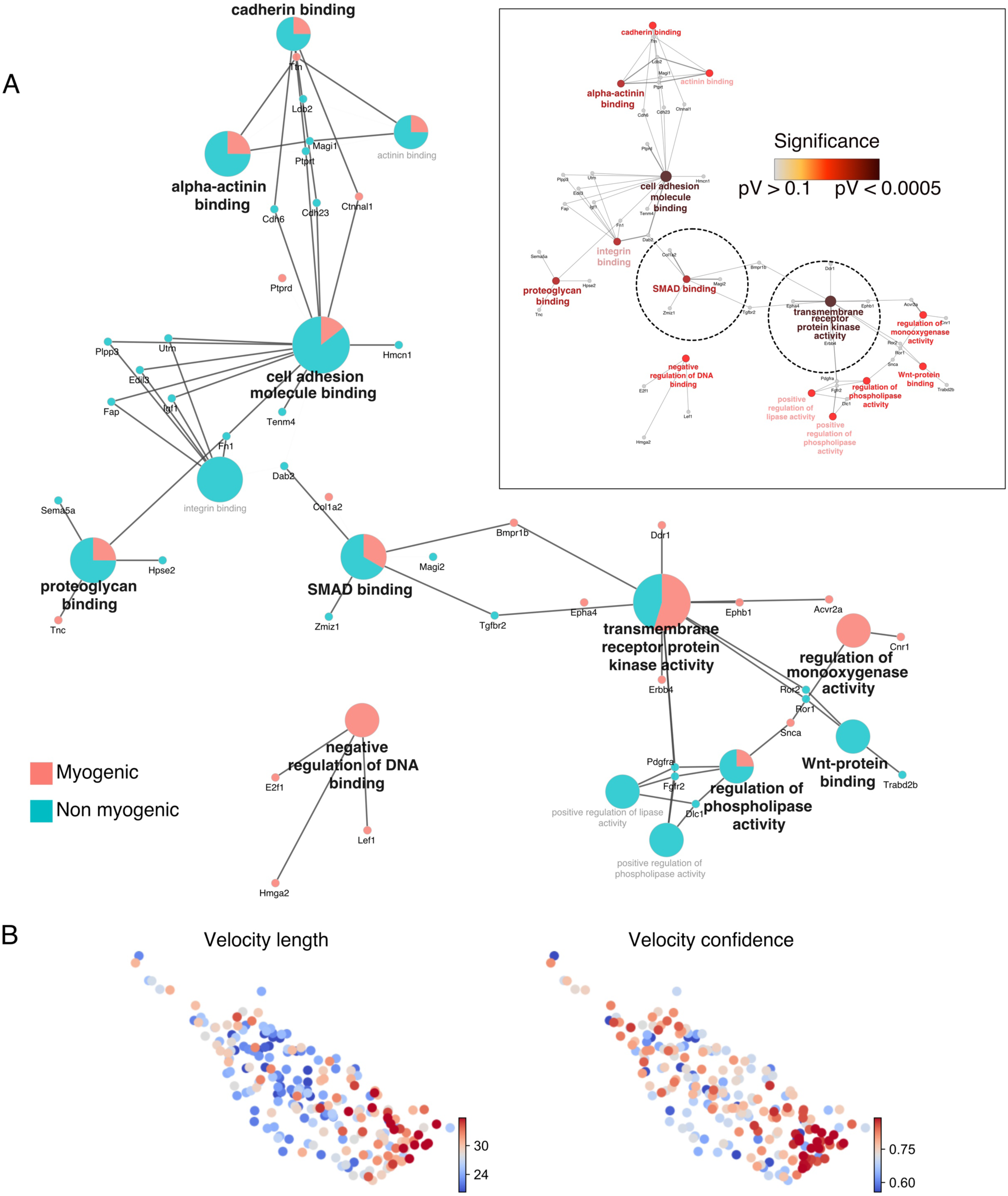
EOM non-myogenic cells arise from a myogenic compartment and crosstalk with myogenic cells. A) GO Molecular Function network, including relative contribution of each cluster to the term and significance levels. Insert show the significance of each term. B) UMAP of *Myf5^Cre/+^; R26^mTmG/+^* E11.5 EOM illustrating velocity confidence and velocity length. Higher confidence is found on both ends of the EOM cluster.

**Supplemental Figure S5.**
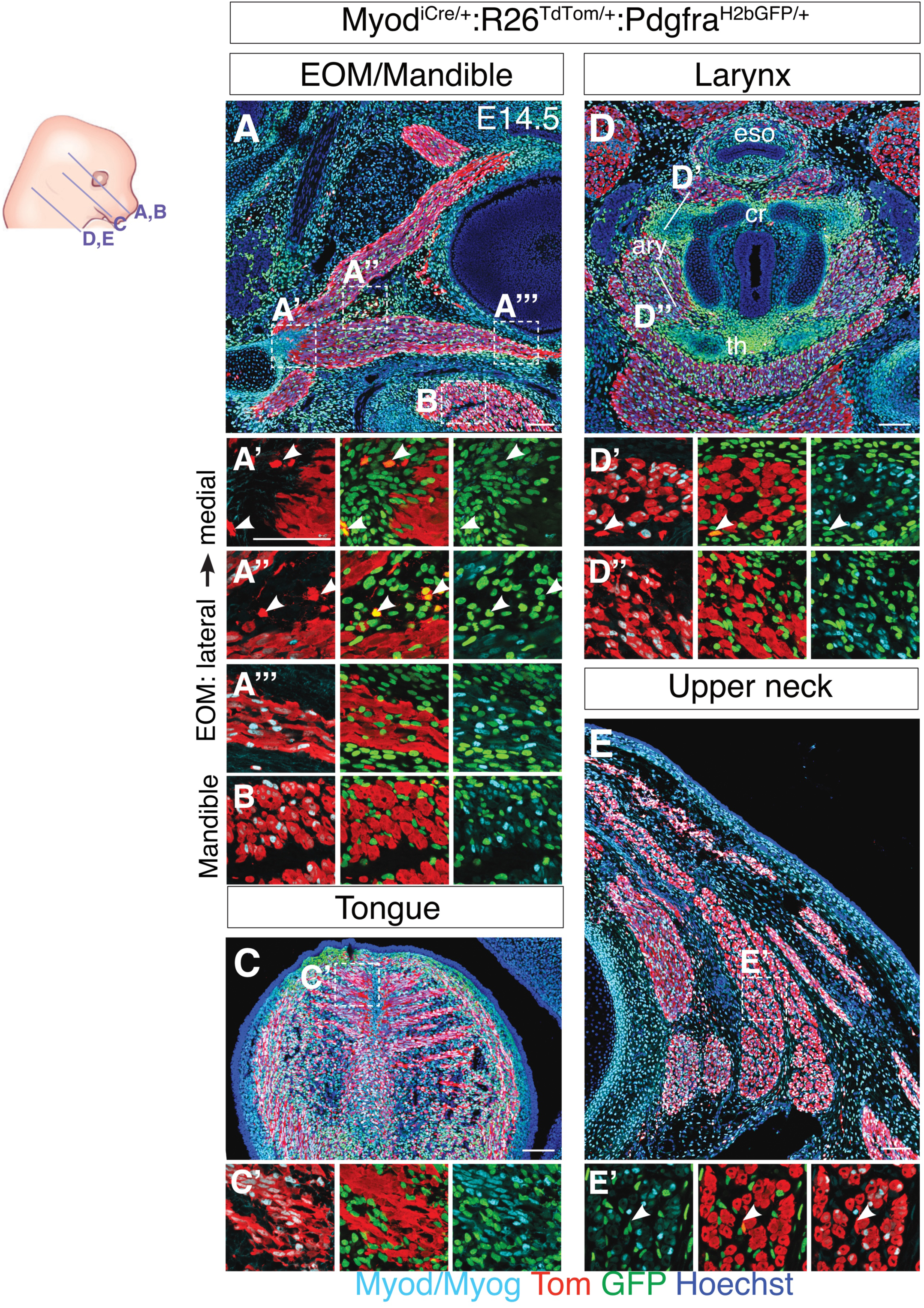
Myod+ cells are restricted to a myogenic fate. A-E) Transverse sections of *Myod^iCre/+^; R26^TdTomato/+^; Pdgfra^H2BGFP/+^* embryos at E14.5 immunostained for Myod/Myog (commited and differentiating myoblasts) in the extraocular (A), mandibular (B), laryngeal (C), tongue (D) and upper neck (E) regions. White arrowhead indicates rare double positive cells (GFP+/Tom+).

**Supplemental Figure S6.**
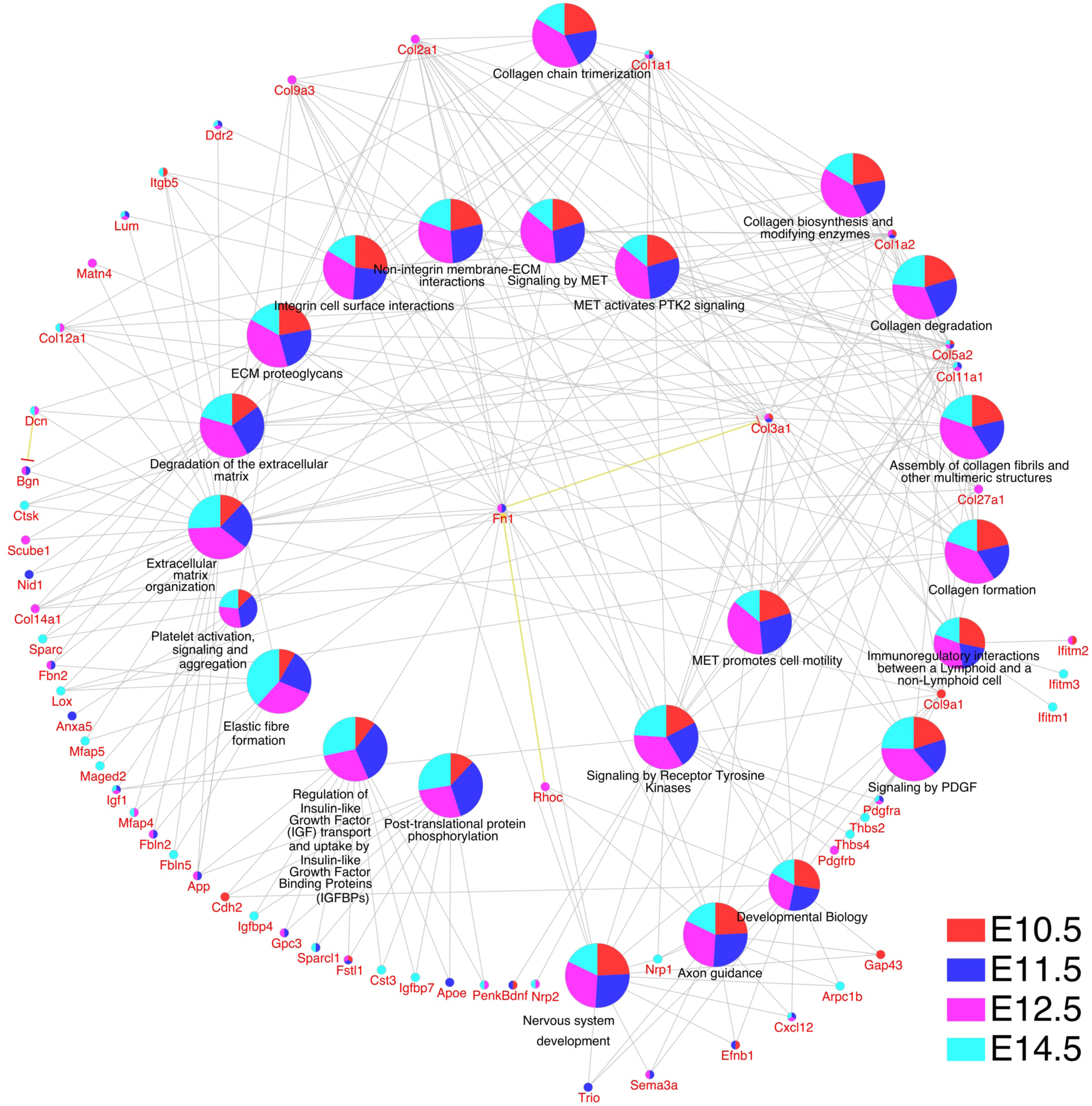
Non-myogenic Myf5-derived cells display a similar gene ontology. Gene ontology analysis for Reactome pathways, including genes underlying each term, and their representation in each dataset. Specific genes of each stage appear related.

**Supplemental Figure S7.**
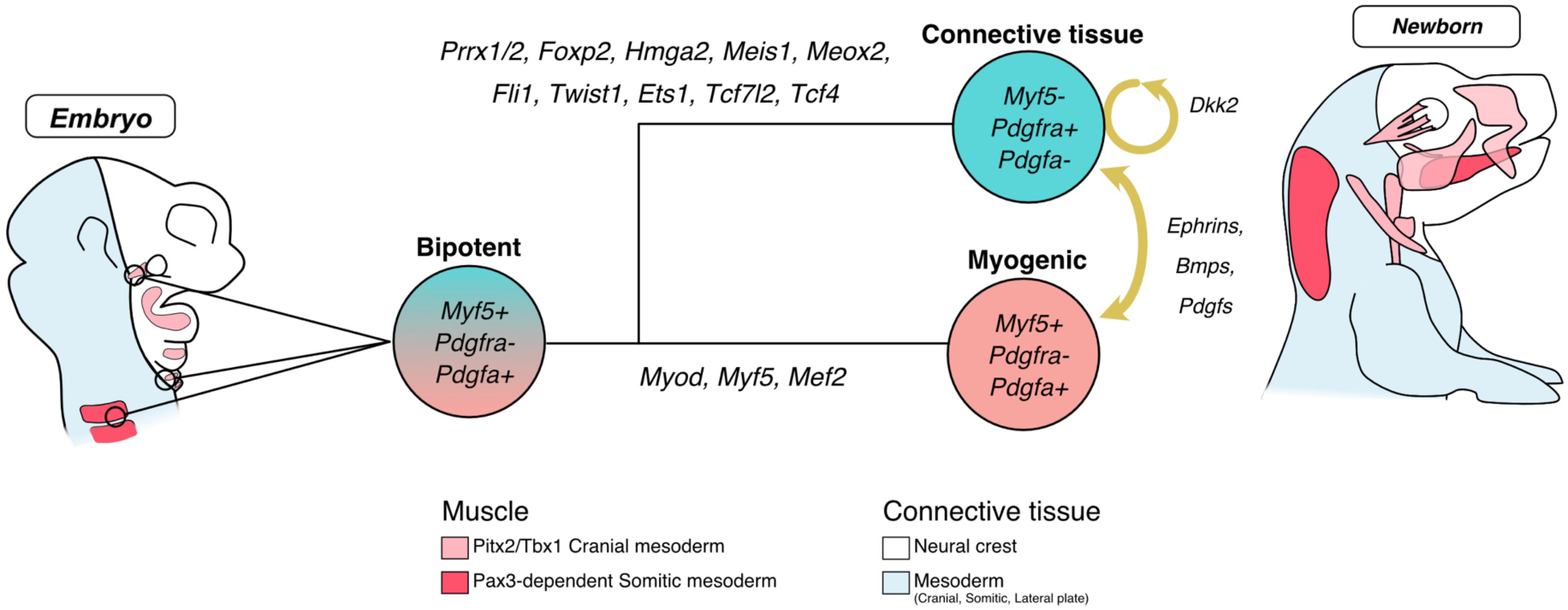
Model of Myf5+ bipotent progenitors giving rise to muscle and associated connective tissues. Model for bipotent Myf5+/Pdgfa+ progenitors giving rise to myogenic and non-myogenic cells and discrete parts of the head, deprived of neural crest. Upon activation of a set of transcription factors including Prrx1/2, Foxp2, Hmga2, Meis1, Meox2, Fli1, Twist1, Ets1, Tcf7l2 and Tcf4, a fibrogenic fate is acquired. A molecular dialogue is initiated at the branchpoint including extracellular matrix components and tyrosine kinase signalling such as Pdgf, Ephrins and Bmps. The non-myogenic fate is maintained cell-autonomously by a canonical Wnt positive feedback loop.

